# A MammaLian Demographic Database for comparative analyses of evolutionary biodemography: malddaba

**DOI:** 10.1101/2025.07.01.662579

**Authors:** V. Ronget, JF Lemaître, B. Spataro, L. Humblot, JM Gaillard

## Abstract

The question of how individual age influences demographic transition rates, such as survival probabilities and reproduction rates, has long been a main question for demographers and evolutionary ecologists. This has resulted in the accumulation of studies estimating the age-specific demographic transition rates across the tree of life over the last few decades. Although this accumulation of studies has enabled comparative demographic analyses to be developed, such analyses remain difficult to perform because age-specific data are scattered across literature. Here, we present a new open-access database, malddaba, which compiles age-specific demographic rates for mammals in the wild from published information. Currently, the database encompasses 170 species from 250 publications, representing 428 age- and sex-specific survival series and 199 age- and sex-specific reproduction series. Each series is reported using a standardized approach aimed at facilitating the extraction and use of that dataset by anyone interested in comparative biology. We show how malddaba can be used to address a variety of questions, ranging from comparative ageing with the assessment of actuarial and reproductive senescence patterns in a wide diversity of mammals. We can also address questions related to comparative population dynamics. For this purpose, in addition to the raw demographic rate records, we were able to build 43 population-specific life tables using malddaba records, which allow demographic outputs to be estimated accurately, such as population growth rate and generation time for those species. The malddaba database will be regularly updated to keep adding new demographic estimates and bring a comprehensive and dynamic view of the diversity of demographic trajectories across mammals.

## Introduction

Demography is the science dedicated to the study of population structure and dynamics. For a long time, demographic analyses have focused on the mortality component following pioneer work designed to describe and understand variation in the duration of life. The first tabulation of the age-specific duration of human life was likely performed by Roman actuaries, who used life tables to estimate how long individuals of a particular age category were expected to live to calculate life insurance products (Trennery, 1926). Later on, Georges-Louis Buffon (1749-1767 in Richard, 1858) and Corbyn Morris (1751) provided detailed life tables and discussed age-specific variation in the duration of life among the Parisians and the Londoners, respectively. The first life table on non-human organisms was published much later by Pearl and Parker (1921) on the normal wild-type *Drosophila melanogaster* and its mutant *‘vestigial’*. However, the demography of a given population is not only based on mortality schedules, but should also include fecundity schedules. Thus, the fundamental demographic equation (also known as the Euler-Lotka equation) Lotka (1907) first enounced in continuous time and based on initial work by Euler (1767) demonstrates that the rate of change in population size between two-time steps is entirely defined by age-specific changes in mortality and fecundity schedules. Lamont Cole (1954) rewrote the fundamental equation of demography in discrete time, opening the door to the scientific field of biodemography, which allows studying population changes in virtually all species over the tree of life. Nowadays, a life table is implicitly a tabulation of the *l_x_* (*i.e.* cumulative survival) and *m_x_*(*i.e.* fertility) series for each age *x* (Williams et al., 2002, see Glossary for a definition of specific terms in italic). From these unitary vital statistics, multiple demographic outputs can be estimated, such as the *net reproductive rate (R0)* or the *generation time (T)* for a specific population (Jones, 2021). Patrick H. Leslie (1945) showed that mortality and fecundity series can also be organized in a population projection matrix, the so-called Leslie matrix can then be used to estimate the asymptotic population growth rate, age distribution, and age-specific reproductive values based on matrix calculus (Caswell, 2001). Both approaches lead to the same demographic outputs when correctly executed (see part III.2 and Box 1 for proof of concepts). These two ways of presenting demographic information have been the focus of previous demographic databases: DatLife for avian and mammalian life tables (Scheuerlein et al., 2017 database not available online anymore), COMADRE (animal demography) and COMPADRE (plant demography) for Leslie matrices (Salguero-Gómez et al., 2015, 2016). Whatever the presentation of the demographic models (i.e. *population projection model (PPM)* using life tables or Leslie matrices), the reliability of the estimate of - for example - the natural rate of population increase strongly relies on the reliability of the mortality and fecundity schedules.

A first and obvious requirement is thus to have accurate information of individual age. This is straightforward when monitoring the fate of entire cohorts from birth to death (corresponding to longitudinal or horizontal life tables), as commonly done in humans (*e.g.* the Human Mortality Database maintained by the Max-Planck Institute for Demographic Research, Barbieri et al., 2015) and only possible under lab conditions for other organisms (*e.g.* Leslie & Ranson, 1940 for a case study on *Microtus agrestis*). When studying populations in the wild, things are markedly different. As a rule, cross-sectional (or transversal) life tables are built from samples collected on dead (*d_x_* series) or alive (*l_x_* series) animals (*e.g.* Deevey, 1947). This approach has been most heavily used between the fifties and the eighties (*e.g.* Spinage, 1972). An abundant literature on assessing age has thus become available (*e.g.* Morris, 1972). However, even in mammals in which tooth eruption and tooth wear patterns are often used, age estimates remain inaccurate, especially for old individuals (*e.g.* Hamlin et al., 2000, but see Lu et al., 2023, for recent developments in age estimations based on DNA methylation patterns). Moreover, imperfect detection (when *l_x_* series are used) and unequal detection among ages (when *d_x_* series are used) often lead to biased estimates of age-specific survival. Capturing individuals as young as possible (ideally at birth), marking them, and monitoring them over their entire life (defining the Capture-Mark-Recapture sampling design, CMR) currently is the gold standard to assess demographic patterns in populations in the wild. Initially aimed to provide reliable estimates of population size in demographically closed populations (*i.e.* with immigration and emigration considered negligible) (Lincoln, 1930; Petersen, 1896), CMR models have included estimates of survival probabilities (Cormack, 1964), first as nuisance parameters to account for so to get unbiased estimates of population size (Jolly, 1965; Seber, 1965), before becoming the most efficient way to assess survival patterns (Lebreton et al., 1992). The number of demographic studies based on long-term population CMR monitorings have consistently increased since the eighties (Clutton-Brock & Sheldon, 2010; Sheldon et al., 2022), providing an efficient way of gathering reliable estimates of age-specific survival probabilities with associated standard error (see Figure 1). This has open the door to intensive analyses of demographic ageing patterns in the wild (reviewed in Nussey et al., 2013).

**Figure 1:**
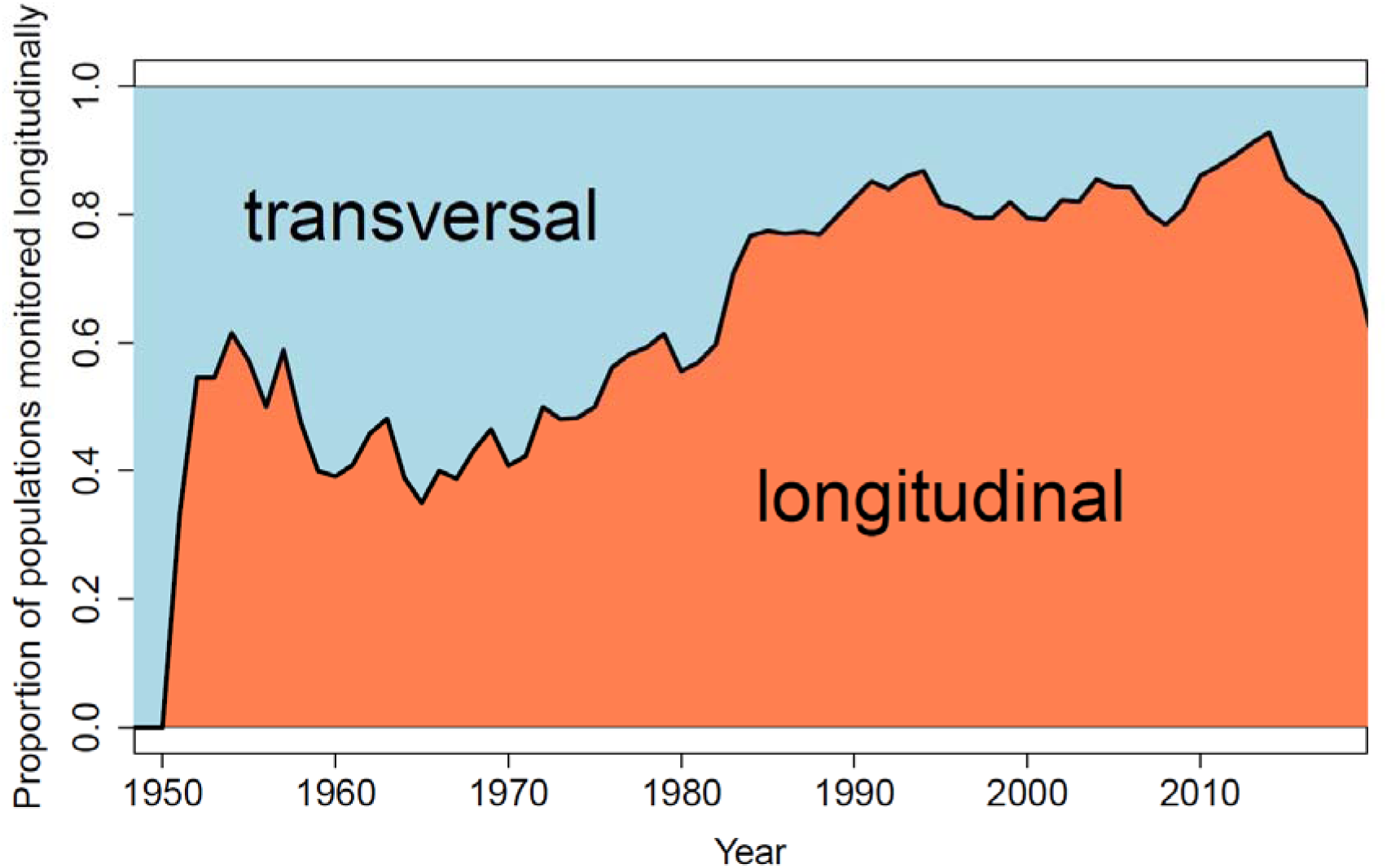
Proportion of populations that are included in malddaba and were monitored longitudinally (*i.e.* using a CMR approach) in orange or transversally (*i.e.* using a cross-sectional approach) in blue. For each population in malddaba, the duration of the monitoring (in years) was determined using the starting and ending date of the monitoring specified in the method section of each paper.

Nowadays, detailed age-specific information of mortality and fecundity is accumulating, which is helpful for comparative biologists to understand the diversity of demographic trajectories across the tree of life (Colchero et al., 2019; Conde et al., 2019; Jones et al., 2014; Reinke et al., 2022). The aim of malddaba (https://malddaba.univ-lyon1.fr/) is thus to compile and report all available information on age-specific demographic data for any population of mammals in the wild or under semi-captive conditions (*i.e.* when individuals can reproduce freely). This unique corpus of data is intended to allow performing thorough analyses of comparative demography to answer a large range of questions in various scientific fields.

## 1) A compilation of a broad range of age-specific demographic traits

The dynamics of any closed population involves only two processes, namely birth and death. To account for the age effect that affects both birth (*i.e.* recruitment that corresponds to birth transitions from all ages to the first age identified in the population census) and death (*i.e.* survival probabilities that correspond to the survival transitions from age *x* to age *x+1*) processes, malddaba reports sex-specific demographic data, associated to age-specific survival probabilities (*S_x_*series) and age-specific fecundity, focusing notably on the reproduction rates (*m_x_* series).

### 1.1) Survival probabilities

The most basic demographic information we need to define the duration of life of a given individual includes its dates of birth and death. Thus, individual lifespans in a population can be computed as the differences between dates of death and dates of birth (Ronget et al., 2024). From a representative sample of individual lifespans or ideally all individual lifespans we can calculate age-specific probabilities of surviving from one time step *x* to the next *x+1*. This conditional survival probability (*i.e.* the probability for an individual to survive to age *x+1* knowing that the individual has already survived to age *x*) is the core information we report in malddaba for survival transition. When individuals are monitored with a perfect detection rate (*i.e.* all individuals alive are consistently controlled at each population census) age-specific survival probabilities can be directly calculated as the ratio of the number of individuals alive at a given census at age *x+1* divided by the number of individuals alive at age *x* in the previous census. However, in most field studies, individuals cannot be controlled at each census, leading detection to be imperfect (Gimenez et al., 2008). Capture-Mark-Recapture (CMR) methods have been developed specifically to tackle this issue by accounting for imperfect detection when estimating survival probabilities (reviewed in Lebreton et al., 1992). The standard approach to estimate survival probabilities consists in modeling both capture probability (*i.e.* the probability to control a given individual at a given census) and survival probability. In addition to report the type of monitoring and methodology associated to each survival series, we also reported the sex associated to each estimate when it was possible. We tried as much as possible to report sex-specific estimates for survival as it is already well-known that there are strong differences in survival between males and females in mammals (Lemaître, Ronget, Tidière, et al., 2020). Estimates for combined sexes were only reported when they were the only data presented in the study.

Long-term *longitudinal* monitoring is now the gold standard, but a substantial number of published studies have (and still) used *transversal* (or *cross-sectional*) data to estimate age-specific survival probabilities through *l_x_* (*i.e.* cumulative probability of surviving from birth to a given age x) or *d_x_* (*i.e.* number of individuals dying between age *x* and age *x+1*) series. While informative, survival analyses from transversal data require additional assumptions such as stationary or stable populations with known population growth rate and thus provide lower quality estimates of survival probabilities compared to survival analyses from *longitudinal* data. Moreover, *transversal* life tables often rely on inaccurate assessment of age (Hamlin et al., 2000), and requires equal detection among individuals across ages, while young individuals are often under-sampled because carcasses of young individuals disappear faster and are harder to find than adults ones (Bodkin et al., 2000). Although age-specific survival probabilities estimated from transversal life tables are less reliable than survival estimates derived for longitudinal studies, they represent a substantial proportion of the populations for which age-specific survival probabilities were retrieved (55% of survival records in malddaba).

### 1.2) Reproduction rates

The second fundamental process driving population dynamics corresponds to recruitment through reproduction. We looked for the standard metric corresponding to the average number of daughters produced alive at birth by females of a specific age (*m_x_* series). Depending on the reproductive seasonality of the species, age-specific fertility can be associated either to a specific age or to a specific age interval. For species experiencing a marked seasonality such as in temperate areas, reproduction occurs each year during a restricted time period within a year (birth pulse species sensu Caughley, 1977). On the other hand, in species experiencing weak seasonality such as in tropical areas, reproduction occurs continuously throughout the entire year (birth flow species sensu Caughley 1977), although in most cases most births take place during a restricted period within the year. We thus defined age-specific fertility (*m_x_*) as the average number of daughters produced by a female of a given exact age *x* for birth pulse species and as the average number of daughters produced by a female of age *x* between age *x* and age *x*+1 for birth flow species.

Despite the strong mother-offspring association existing in mammals in response to the need of maternal care during the lactation period, it is typically difficult to assess accurately the reproductive status of a given female at a given age. The reproduction process can be considered as a sequential process (see Lemaître & Gaillard, 2017) including successively mating competition, fecundation, pregnancy, parturition and lactation. Most studies have focused on partial information associated with one or more components of the reproductive process, usually the easiest to monitor in the focal species. We thus did not only include *m_x_* series in malddaba but retrieved information from all traits related to reproduction. This encompasses prenatal metrics of reproduction such as pregnancy rate or litter size *in utero*, metrics at birth such as *m_x_*, proportion of parturient females, or litter size, and postnatal metrics such as proportion of females with offspring, litter size at weaning, or first year survival of offspring. While the combination of those different metrics can allow computing *m_x_*it was not always the case. Due to the plurality of reproductive metrics, comparative analyses of age-specific reproduction are much scarcer than comparative analyses of survival (but see Lemaître et al., 2020). Similar to survival, age-specific reproductive data compiled in malddaba are sex-specific. Indeed while biodemography research has for a long time focused on females due to their central role in population dynamics (Archer et al., 2022), age-specific reproductive data in males are now commonly reported (Cambreling et al., in press) thanks to the increasing use of paternity analyses in the study of animal populations. Overall, these data encompass a wide range of metrics describing both mating success (e.g. probability of mating) and reproductive success (e.g. number of sired offspring, probability of paternity), those male-specific data on reproduction will be added in malddaba in the near future.

Most data collected on reproductive traits come from longitudinal studies although some metrics have been calculated based on transversal data (45% of populations with reproductive traits obtained from transversal data). For instance, shot individuals of game species are routinely used to measure the proportion of pregnant females as well as the number of embryos per breeding female. As for survival data, reproductive traits estimated from transversal studies were considered of lower quality for much of the same reasons (inaccuracy in age assessment and potential sampling biases caused by unequal susceptibility to hunt by breeding vs. non-breeding females).

## 2) The database

### 2.1) Data inclusion

The malddaba database encompasses data collected at the population level and coming from papers published in journals, books, preprints, theses or technical reports. Most of them come from previous partial compilation of studies done by the authors (e.g. in Lemaître, Ronget, & Gaillard, 2020; Lemaître, Ronget, Tidière, et al., 2020) in addition to specific literature research using Google Scholar and Web of Science databases. Our two main inclusion criteria to include demographic estimates in malddaba were first to originate from non-captive populations and second to be published. Nonpublished data were only included if those estimates are associated with a population in which some demographic estimates were already published and for which metadata are available. In that case, nonpublished data are only compiled in malddaba with the authorization of the owners.

### 2.2) Data extraction

In addition to reporting age-specific demographic estimates, malddaba records information from the initial publication used to assess the monitoring and methodology as well as the species studied. Collected data is organized in a SQL model implemented with PostgreSQL. We classified this information into six different categories: “Data table”, “Demographic trait”, “Species”, “Location”, “Reference”, and “Study”. In the database, each record (or line) is a unique combination of the specific demographic trait, the sex, the species, the location and the publication. For example, a study where survival data for both sexes and reproduction data for females (e.g. litter size and *m_x_*) are available, will be associated with 4 different records in malddaba: one record for age-specific female survival, one record for age-specific male survival, one record for age-specific female litter size, and one record for the female *m_x_* series. All extraction procedures associated with the different fields (or columns) of each category will be explained hereafter. A summary of the column names in the database associated to each category is presented in Table 1.

**Table 1:** Information reported for a given study included in malddaba (See data extraction in text for more details about the extraction procedure)

| Category | Field name | Description |
| --- | --- | --- |
| Species |  | All information associated with the taxonomic name of the species monitored. All names are standardized according to the Catalogue Of Life nomenclature |
|  | Genus | Genus name (latin) |
|  | Species | Species name (latin) |
|  | Sub.species | Sub-species name (latin) |
| Location |  | All geographical information to locate the monitoring site |
|  | Study_site | Name of the study site as reported in the publication |
|  | Country | Name of the country in which the study took place |
|  | GPS_coordinates | GPS coordinates of the study sites in hexadecimal format |
| Demographic Trait |  | Information on the type of demographic trait reported |
|  | Trait_type | Category of demographic data reported: either “survival” or “reproduction” |
|  | Trait | Specific demographic trait reported<br>- “Survival_rate” or “N_dead” or “N_alive” for “survival” type<br>- “P_reproduce” or “Number of offspring” or “Litter_size” or “mx” or “Juvenile_survival” for “reproduction” type<br>(See text for more information on the meaning of each trait) |
| Reference |  | Information about the reference used to extract the data |
|  | DOI | DOI for the publication when available |
|  | APA | Full citation of the reference used in APA style |
| Study |  | Information about the sampling of individuals in the study and |
|  |  | the methodology used to estimate the demographic traits |
|  | Monitoring_type | Type of monitoring: “longitudinal” or “transversal_dx” or “transversal_lx”<br>(See text for more information on the meaning of each category) |
|  | Sex | Sex of the individuals: “Females”, “Males” or “Combined” |
|  | Number_of_individuals | Number of unique individuals sampled for building the dataset of the focal study (different from the number of captures because an individual can be captured multiple times in a longitudinal study) |
|  | Number_of_captures | Number of total captures for building the dataset of the focal study |
|  | Start_year | Year when the monitoring started |
|  | Duration | Duration of the monitoring in years (only for data included in the specific record) |
|  | Right_censored | Whether any individual in the focal study was right-censored (“Y”) or not (“N”)<br>When individuals are right-censored some individuals were still alive after the last age reported in the focal study, but those ages were not included in the demographic estimates reported |
|  | Primary_data_extracted | Whether raw estimates of demographic data are reported (“Y”) or not (“N”). Estimates from full age-dependent CMR models are considered as raw estimates |
|  | Modeled_data_extracted | Whether estimates from continuous age models are reported (“Y”) or not (“N”) |
|  | Model_description | Description of the continuous age-dependent model used to compute the age-specific demographic estimates. Only reported if Modeled_data_extracted is “Y” |
|  | Reproduction_time | Time at which the reproductive traits were measured, only |
|  |  | reported for reproductive data |
|  | Birth_pulse | Information about the reproduction timing within a specific age class:<br>- if “Y”, the focal species displays a birth pulse type so that all reproduction consistently takes place at the starting age of any age interval<br>- if “N”, the focal species displays a birth flow type so that all reproduction takes place between the starting and the ending age of any age interval |
|  | Graph_extraction | Whether demographic estimates were extracted from graphs (“Y”) or not (“N”) |
|  | Comments | Any comment associated with the extraction of the data |
|  | ID | Specific unique ID of the record in malddaba |
| Data_table |  | Information on the age-specific values of the demographic estimates for the specific record |
|  | Life_table_id | Specific unique ID of the record in malddaba<br>(used to connect “metatable.csv ” and “lifetable.csv” tables only displayed if the users are extracting database information using the “csv” format files) |
|  | start_age | Age at the start of the specific age-class in years |
|  | end_age | Age at the end of the specific age-class in years |
|  | Mean_parameter | Raw demographic estimates associated with the specific age-class, only reported if Primary_data_extracted is “Y” |
|  | Modeled_mean_parameter | Demographic estimates from continuous age-dependent models associated with the specific age-class, only reported if Modeled_data_extracted is “Y” |
|  | N_alive | Number of individuals alive at the start of the age-class if |
|  |  | Trait_type is “survival”, age-specific sample size otherwise |
|  | N_dead | Number of individuals dying during the specific age-class |

#### 2.2.1) Data table

In this category, age-specific demographic estimates are reported in the same format for survival and reproduction data.

##### Survival data

For survival data, information was reported differently whether the survival estimates were computed from a longitudinal or a transversal monitoring. For longitudinal data, survival probabilities from a starting age (“start_age”) to an ending age (“end_age”) in years were reported. In the case of estimates from CMR studies, survival probabilities from full age-dependent models were reported in the “mean_parameter” column (i.e. when age-specific survival probabilities are modelled independently). In addition, the number of individuals at risk of dying (*i.e.* number of individuals alive at the beginning of the starting age) was reported when available in the “N_alive” column. The number of individuals dying in the specific age interval was also reported in the “N_dead” column when available. For transversal data, we prioritized reporting only either the number of individuals alive at the beginning of each age class for *transversal l_x_* data in the “N_alive” column and the number of individuals found dead during the specific age class in the “N_dead” column for *transversal d_x_* data.

In some studies, survival probabilities were estimated using a continuous age-dependent model. In that case, survival estimates from this model were reported in the “Modeled_mean_parameter” column. All information regarding the type of continuous model fitted is reported in the “Model_description” column.

##### Reproductive data

We reported each estimate of a reproductive parameter in the “mean_pararameter” column as well as the sample size associated for each age in the “N_alive” column when available. We presented reproduction estimates into five main trait categories: (1) the probability to reproduce defined as the probability for all individuals of a given age to be pregnant (for females), to give birth, or to have successfully raised at least one offspring depending of the timing of the sampling, (2) the litter size defined as the number of offspring produced during pregnancy (*i.e.* number of corpora lutea or number of embryos, for females only), at birth (*i.e.* number of newborns), or during the dependent period of the offspring (*i.e.* number of offspring that survived for a while after birth), (3) the number of offspring produced by all females or males of a given age (*i.e.* pooling both successful and unsuccessful females), (4) the *m_x_* defined as the average number of daughters produced alive at birth by a given individual of a given age. Approximations of *m_x_* series were used in some studies where offspring cannot be observed right after birth. For instance, in species with offspring born in a den, *m_x_* was measured at emergence. Likewise, mating success is more often reported than birth rate in males so that male *m_x_* was generally measured at mating rather than at birth, and (5) juvenile survival defined as the offspring survival probability of females or males of a given age from birth to a specific young age. Juvenile survival was often calculated over a different time interval, using either a specific time period after birth in weeks or months or a specific life history event such as offspring survival until weaning or independence.

We informed whether the reproduction was with a birth pulse or a birth flow type in the “Birth_pulse” column. In case of a birth pulse species, reproduction was assumed to take place at the start of the age interval (“start_age” column), whereas for a birth flow species, the reproductive event was assumed to take place continuously during the age interval (from “start_age” to “end_age” columns). For both reproduction types the value of the reproductive trait corresponded to the total reproduction over that interval. In addition, we reported the timing of the measure in the “reproduction_time” column (i.e. whether the reproductive trait was measured at pregnancy, at birth, or after birth). For instance, a pregnancy probability corresponds to a probability to reproduce measured at pregnancy in the Maldabba notation.

Like survival, reproduction was sometimes estimated using a continuous age-dependent model. Those modelled values were reported in the “Modeled_mean_parameter” column. All information regarding the type of continuous model fitted was also reported in the “Model_description” column.

#### 2.2.2) Demographic trait

In this category all information to identify the demographic trait studied is reported. It allows the user to know whether it is a survival or a reproduction record in the “Trait_type” column and which specific trait is reported in the “Trait” column (“Survival_rate” or “N_dead” or “N_alive” for survival data, or “P_reproduce” or “Number of offspring” or “Litter_size” or “mx” or “Juvenile_survival” for reproduction data).

#### 2.2.3) Species

For each record, information on the name of the genus (“Genus” column), species (“Species” column) and subspecies (“Sub.species” column) was reported following the catalogue of life (COL) nomenclature (https://www.catalogueoflife.org). Older species names from original publication were changed to match the actual COL nomenclature.

#### 2.2.4) Location

For each record, information to locate the study site was reported. We reported the name of the study site (“Study_site” colum), the name of the country (“Country” column) as well as the geographic coordinates (in hexadecimal format) of the specific study site (“GPS_coordinates” column).

#### 2.2.5) Reference

As we aim our extraction procedure to be as transparent and reproducible as possible all reference information associated with the original publication was reported. DOI (“DOI” column) if available and full citation in APA style were compiled (“APA” column).

#### 2.2.6) Study

General information associated with the study was reported in malddaba. Information on the sex under study (“Sex” column), whether total sample size was reported as either the total number of unique individuals included in the study (“Number_of_individuals” column) or the total number of captures used to compute the demographic estimates (“Number of captures” column), the type of monitoring (longitudinal vs. transversal) (“Monitoring type” column), of the time period the study covered (“Start_year” and “Duration” column), and whether there was any right censoring in the data (*i.e.* when information on oldest individuals was excluded or absent, “Right_censored” column) were reported. We also added any information relevant about the extraction procedure used (See Table 1 for a detailed description of all fields).

### 2.3) Database accessibility

All the demographic estimates and the information associated are freely available to download on the website of the database (https://malddaba.univ-lyon1.fr/). Demographic estimates can be selected and downloaded in different format over the tab “DATABASE” (https://malddaba.univ-lyon1.fr/pages/database.php). It is possible to look at specific records using the web interface or directly extract and download the records in the database using the “export row” column. Users can either download all the database using the select all functions or use the search function to look for specific type of demographic estimates or species. Data can be downloaded in 3 different formats: csv files, Rdata list or JSON array. For each file type a compressed zip file is downloaded with the specific data file and a “README.txt” file with the explanation relative to each column from the data file (see Table 1). As we strive to have a comprehensive view of all age-specific demographic estimates in mammals in the wild, we encourage researchers with demographic data on mammals that are not already reported in malddaba to reach us using the contact page in the website (https://malddaba.univ-lyon1.fr/pages/contact_us.php).

### 2.4) Overview of malddaba

On 10/06/2025, malddaba compiles demographic estimates from 250 unique publications. The oldest paper included in the database was published in 1956. As expected, the literature corpus mostly includes articles about demography, population dynamics, life-history, and senescence (See Fig 2A). Populations studied were located all around the world but with a large majority studied in three main areas: Europe, North America, and Sub-Saharan Africa (See Fig 2B). The malddaba database includes a large diversity of species with 171 species associated with 428 survival records and 199 reproduction records. While most mammalian taxa are represented, there is a disproportionate importance of *Artiodactyla*, *Carnivora* and *Sciuridae* that are intensively studied. The database will be updated regularly to add new species and include survival or reproductive information when missing. Note that for most species data are only associated with one population. We also aim to increase the number of populations of a given species in the future to allow the user to assess between-population variation in demographic estimates as well as between-species variation. For instance, the species for which we currently have the highest number of populations is *Trichosurus vulpecula* with 16 records from 9 different populations.

**Figure 2:**
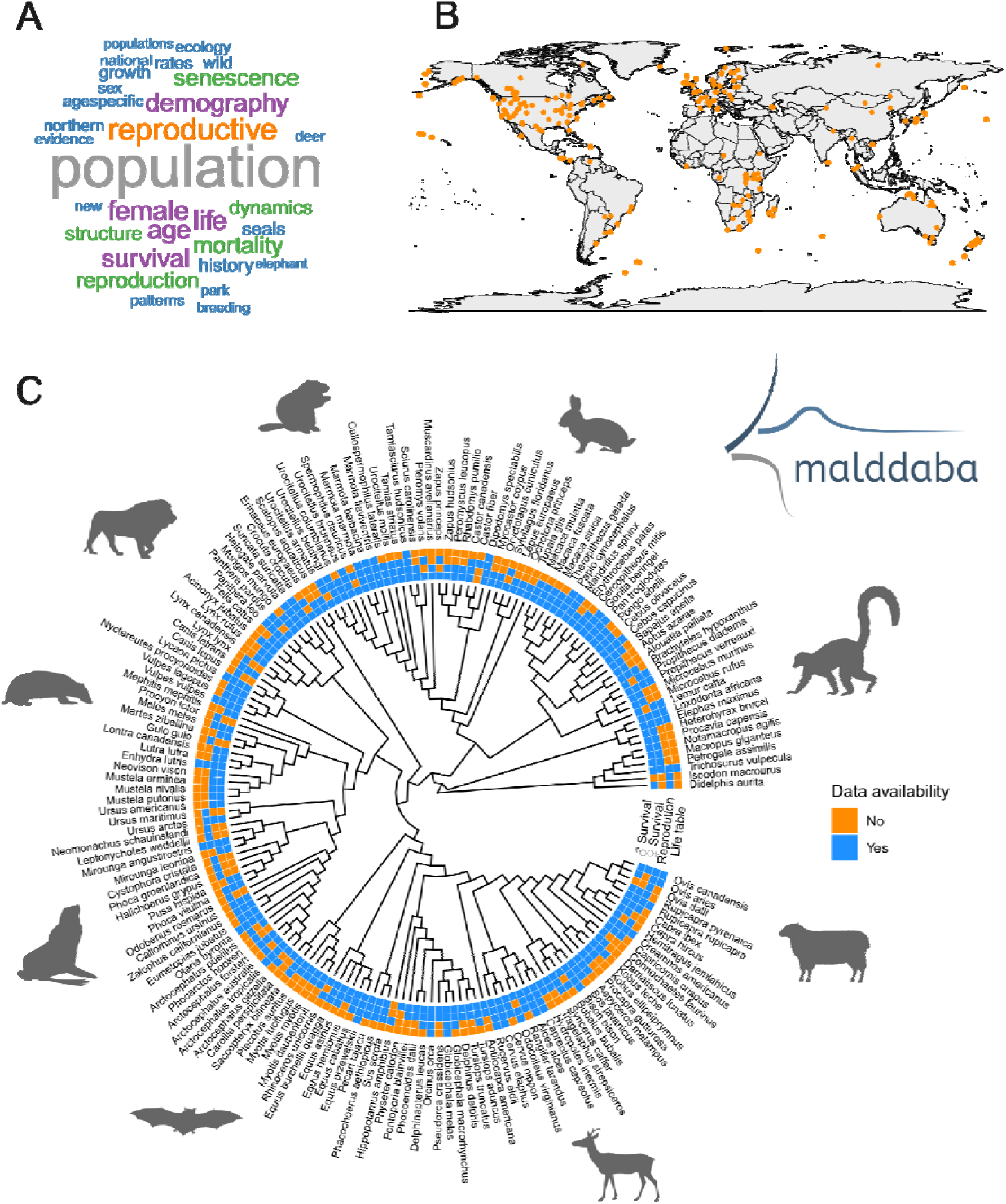
(A) Word cloud associated with the title of all the 250 publications included in malddaba: the bigger the word, the more likely the word is present in the title. (B) World map of all the different locations included in malddaba (orange dots). (C) Phylogeny of all the species included in malddaba. Each colored square represents whether at least one demographic record is available for male survival, female survival, reproduction, or life table data (i.e. full age-dependent survival and reproduction available for the same population) in blue from inner to outer circles, respectively. The phylogeny was built using the “rotl” package (Michonneau et al., 2016).

## 3) Examples of research questions that can be tackled using malddaba

We illustrate here several research questions that can be tackled using malddaba. We will first focus on the benefits that malddaba could bring to the comparative biology of ageing research field.

### 3.1) The relevance of malddaba for multi-disciplinary ageing research

One of the most variable life history trait in the living world is lifespan (Carey & Judge, 2000) and researchers are currently taking advantage of this huge variation to identify private or evolutionary conserved mechanisms that could be associated with between-species differences in lifespan (e.g. Haghani et al., 2023; Tyshkovskiy et al., 2023 for some recent examples). Many of these groundbreaking studies are focusing on taxonomic groups displaying a so-called ‘exceptional lifespan’ (*e.g.* species supposed to display particularly long life for their body mass according to allometric rules, but see Gaillard & Yoccoz, 2024 for alternative definitions of exceptional lifespan) as they can bring insights on fine-scale physiological and molecular pathways conferring protection against common causes of death, such as infectious diseases or cancer (*e.g.* Huang et al., 2019 in bats or Gorbunova et al., 2014 in rodents). Although these studies have been successful in identifying relevant mechanisms (Austad, 2022), we emphasize that age-specific mortality data compiled in malddaba constitute a powerful tool to push the field further, by removing the inherent methodological and biological limitations to the use of lifespan metrics, and maximum lifespan in particular. There is indeed compiling evidence that maximum lifespan is only a poor descriptor of a species lifespan (Ronget & Gaillard, 2020) and that the use of this metric can lead to spurious conclusions (*e.g.* Rozing et al., 2017 in the specific context of lifespan limits in humans). Maximum lifespan is known to be extremely sensitive to sample size (Krementz et al., 1989) and environmental conditions (Lemaître et al., 2014). For instance, the comparison of two ‘maximum lifespan databases’ has revealed that the estimation of evolutionary allometric exponents (*i.e.* parameters that characterize the increase in lifespan as a function of body mass across species) markedly differ between these distinct data sources (Lemaître et al., 2014). This is likely explained by the relative proportion of records retrieved from captive populations between the two databases, a factor that is known to sometimes lead to unrealistic maximum lifespan. Moreover, this heterogeneity in the data sources complexifies the identification of species displaying exceptional lifespan, a topic that already suffers from poor methodological approaches (see Gaillard & Yoccoz, 2024 for a recent review). In that context, malddaba offers new perspectives for comparative analyses aiming to focus on lifespan by providing raw data on age-specific mortality that enables to quantify much more biologically relevant metrics of lifespan (*i.e.* unbiased by extreme values) as well as their associated error measurements. For instance, adult lifespan measurements such as the adult lifespan 80% or 90% (*i.e.* when 80% or 90% of the individuals alive at the onset of adulthood are dead), which encompass most of the old age classes without being biased by the few particularly long-lived individuals, constitute much more accurate metrics (Moorad et al., 2012) when it comes to identify the ecological and biological drivers of lifespan (Lemaître, Ronget, Tidière, et al., 2020).

Moreover, one fundamental point to bear in mind when performing comparative analyses on ageing is that lifespan metrics (whatever the statistical approach used to estimate them) do not measure only the change in age-specific mortality risk throughout life. Indeed, while common beliefs sometimes assume that the rate of actuarial senescence and lifespan are exchangeable metrics conveying similar information, this is in fact far to be true (Kowald, 2002; Ronget & Gaillard, 2020). For instance, using demographic data on mammals, Péron and colleagues (2019) have demonstrated that the variation in actuarial senescence rates explains less than half the variance in lifespan. Lifespan, which by definition ‘simply’ corresponds to the duration of life (Ronget et al., 2024), is thus irrelevant to describe the complexity of the age-specific decline in survival probabilities with increasing age. Interestingly, biomedical studies have started to document the complexity of the age-specific changes in the molecular and physiological pathways likely involved in the ageing process (Lehallier et al., 2019) and there is now a need to identify the (epi-)genetic and physiological pathways underlying the age at the onset of actuarial senescence, the rate of actuarial senescence (Lemaitre, Gaillard, Pavard, et al., 2024), as well as explaining the variability in the pace and the shape of ageing among species (see Baudisch, 2011; Ronget & Gaillard, 2020 for further discussion).

Such research objective could now be reached using mammals as a biological model thanks to malddaba. Indeed, mathematical functions (*e.g.* bathtub models such as the Siler model, Siler, 1979) enabling the estimation of key actuarial senescence parameters, as well as their associated standard errors can be easily fitted following the extraction of the age-specific survival probabilities (see Section I), while controlling for methodological factors known to influence ageing parameters (*e.g.* hunting status, Koons et al., 2014) and already available in the current version of the database. So far, the full potential of actuarial senescence metrics in comparative biology of ageing has already been demonstrated in studies seeking to decipher the complex differences between male and female mortality patterns across species (Bronikowski et al., 2022; Lemaître et al., 2024) and notably through the use of malddaba (Lemaître, Ronget, Tidière, et al., 2020).

Overall, malddaba offers promising opportunities for the identification of mechanisms shaping the changes in age-specific mortality risk and associated with species displaying a particularly late onset or low rate of actuarial senescence. One may argue that the number of mammalian species for which data on maximum lifespan is available is much higher than the number of mammalian species for which age-specific survival data are available, hence potentially diminishing the relevance of malddaba for such studies. However, the set of species already compiled for which actuarial senescence patterns can be estimated is (at the time of the publication of the database) already high (155 species with survival estimates available), encompassing species displaying a high diversity of life history strategies (see section II.4, Figure 2). Moreover, a major asset of malddaba is that it is a population-level database that enables us to perform comparative analyses while simultaneously controlling for environmental effects and increasing the robustness of the outcomes.

Finally, it is important to emphasize that malddaba can be equally relevant for both quantifying and identifying the eco-evolutionary drivers of reproductive senescence. Indeed, the decline in age-specific reproductive performance with increasing age has fundamental implications for eco-evolutionary processes through its impact on fitness (Bouwhuis et al., 2012; Kowald & Kirkwood, 2015) and is tightly associated with reproductive health in both sexes (Comizzoli & Ottinger, 2021). This is thus not a surprise that the search for the drivers of reproductive ageing has recently become of topical importance (Lemaître & Gaillard, 2023). In that context, the spate of age-specific reproductive data compiled in malddaba (and more specifically data on *m_x_* values) have already demonstrated that reproductive senescence metrics covary with the species position along the slow-fast continuum (Lemaître, Ronget, & Gaillard, 2020 for females) and are heavily influenced by female reproductive behaviour (Lemaître, Ronget, & Gaillard, 2020). We now advocate for a much wider use of these data in the context of reproductive ageing, notably to identify, as suggested for actuarial senescence (see above), the mechanisms underlying the age-specific changes in reproductive performance through life, but also the emergence of post-reproductive lifespan across the tree of life.

### 3.2) Demographic analyses

The malddaba database is of great relevance for addressing any question related to population dynamics and comparative demography, in a sex- and population-specific way. This database does not only include data about the adult stage but aims to describe the full trajectory (*i.e.* from birth to death) of females and males across the life cycle of the focal species. Demographic transitions in terms of survival and reproduction are required to describe the whole life cycle of a given species and thereby to estimate the demographic outputs at the population level. *Population Projections Models* (PPM) have been devised with that aim to specifically track how the age distribution changes from one time step to the next one. Discrete life table analyses and matrix projection population models (MPPM) are typically two different descriptions of the same demographic approach that can be used to project age distributions. Both technics should provide the same demographic outputs when correctly formulated. As this equivalence is not that easy to grasp for non-demographers, which has led to wrong demographic assessments in a substantial number of cases (Kendall et al., 2019), we demonstrate this equivalence in Box 1.

#### Box 1

Correspondence between life table analysis and matrix projection models

Let first consider a population in which survival transitions (*S_x_* series) and reproduction transitions (*m_x_*series) are available. Note that we will only consider discrete models with a birth pulse meaning that all reproduction takes place at a given age and not between two ages. Those transitions can be presented in a life table format (Jones, 2021) by recursively calculating the cumulative survival at a given age (*i.e.* the *l_x_* series) from the *S_x_* series:

| $x$ (Age) | $S_x$ | $l_x$ | $m_x$ |
| --- | --- | --- | --- |
| 0 | $S_0$ | $l_0=1$ | 0 |
| 1 | $S_1$ | $l_1=l_0S_0$ | $m_1$ |
| 2 | $S_2$ | $l_2=l_1S_1$ | $m_2$ |
| 3 | $S_3$ | $l_3=l_2S_2$ | $m_3$ |
| 4 | 0 | $l_4=l_3S_3$ | $m_4$ |
| 5 | | $l_4=0$ | |

Based on that life table with specific *l_x_* and *m_x_* series, different demographic metrics can be calculated such as population growth rate *r* by solving the Euler-Lotka equation of demography, the net reproductive rate *R0*, or the generation time *T* (See supplementary S1 displaying a case study for which all these metrics have been calculated).

From this life table, it is also possible to build the associated matrix population projection models (MPPM) (Caswell, 2000). Each entry of the projection matrix *A_x,x+1_* corresponds to the transition rate from age *x* to *x+1*. The MPPM can be built differently depending on when the population census takes place relative to births (i.e. pre-breeding vs. post-breeding census). Here are the two MPPM for the life cycle used above to present the life table.

Pre-breeding census:

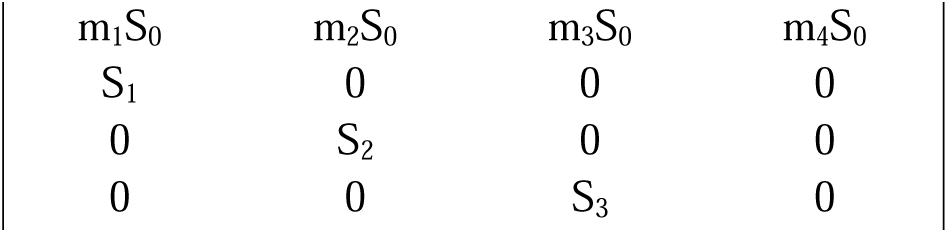

Post-breeding census:

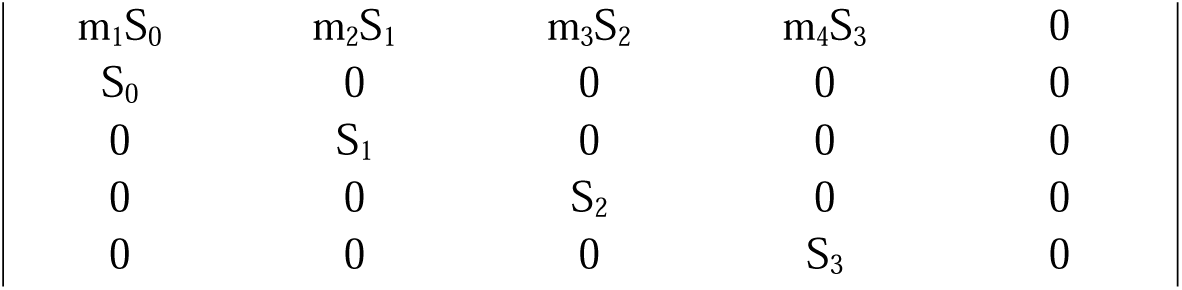

|  |  |  |  |  |  |
| --- | --- | --- | --- | --- | --- |
| | $m_1 S_0$ | $m_2 S_1$ | $m_3 S_2$ | $m_4 S_3$ | 0 |
| | $S_0$ | 0 | 0 | 0 | 0 |
| | 0 | $S_1$ | 0 | 0 | 0 |
| | 0 | 0 | $S_2$ | 0 | 0 |
| | 0 | 0 | 0 | $S_3$ | 0 |

Using matrix calculus, it is possible to compute all demographic outputs such as the asymptotic population growth rate (as the dominant eigenvalue of the matrix), age-specific distribution and age-specific reproductive values (as the right and left eigenvectors, respectively). Both formulations, either with life table and projection matrices, will lead to the same demographic outputs but those dependent on the population age structure because the age distributions sampled during pre- and post-breeding censuses are different.

In addition to reporting all the raw demographic estimates, we built a life table dataset in malddaba by including both reproduction and survival schedules for all species when complete demographic data were available. Using such life tables, demographers can build any population projection model they need and then extract a range of demographic metrics for several species that can be used directly for comparative demographic analyses.

As for the extraction of the main malddaba data, we aimed for maximal transparency and reproducibility for the methodology used to build those life tables. Our two main criteria to build a life table from malddaba records was first to have full age-specific survival and reproduction estimates for the same species and the same population. As transition rates could vary widely among populations, we recommend to build a life table for a specific population only, to avoid any issue associated with mixing transition rates from different populations (Rosa et al., 2025). Second it was only possible to build the life table if survival and reproduction were reported from birth to the oldest ages. For some of the survival records, survival probability for the last age was not exactly 0 meaning that some individuals were still alive/censored at old ages. In those cases, we used the *l_x_* series to assess whether the proportion of censored individuals was high or not. When the proportion of remaining individuals was lower than 5% at the last age, we included the population in the life table dataset by ending the survival series using a survival of 0 at the next age. The importance of those very old individuals is likely negligible for the whole population dynamics, but it is possible to remove those life tables as all decisions associated to the life table building is reported in the dataset.

For each life table, *S_x_* and *l_x_* series are reported as well as *m_x_* series at regular time intervals. The origin of the raw data used to build the life table is reported explicitly and linked to the original malddaba data. Any decision on how to censor or construct the survival series is also reported (see Supplementary S2 for a full description of the life table dataset). Similarly, *m_x_*was calculated based on several other reproductive traits for most populations. The origin of the data, the procedure and the formulas used in that case are all reported in the full dataset (see Supplementary S2). In addition to present the full life table for each population with required data, we also computed common demographic metrics such as the life expectancy, *e_x_*, at a specific age, the asymptotic population growth rate, *r*, the net reproductive rate, *R0*, the reproductive values at each age, *V_x_*, and two measures of generation time T_b_ and T_c_ (see supplementary S2 for the formulas used to calculate each metric).

The life table dataset includes so far 43 species and will be updated as more demographic data in malddaba will be extracted. The dataset is available to download on the malddaba website (https://malddaba.univ-lyon1.fr/pages/life_table_data.php). Those species cover a large diversity of life history strategies from fast-living species with short generation time to slow-living species with long generation time (see Figure 3). While our set of species is quite restricted compared to the full malddaba, we chose to have restrictive inclusion criterion to only get high quality life tables. This dataset could be thus considered as the gold standard to perform any demographic analysis across mammal species.

**Figure 3:**
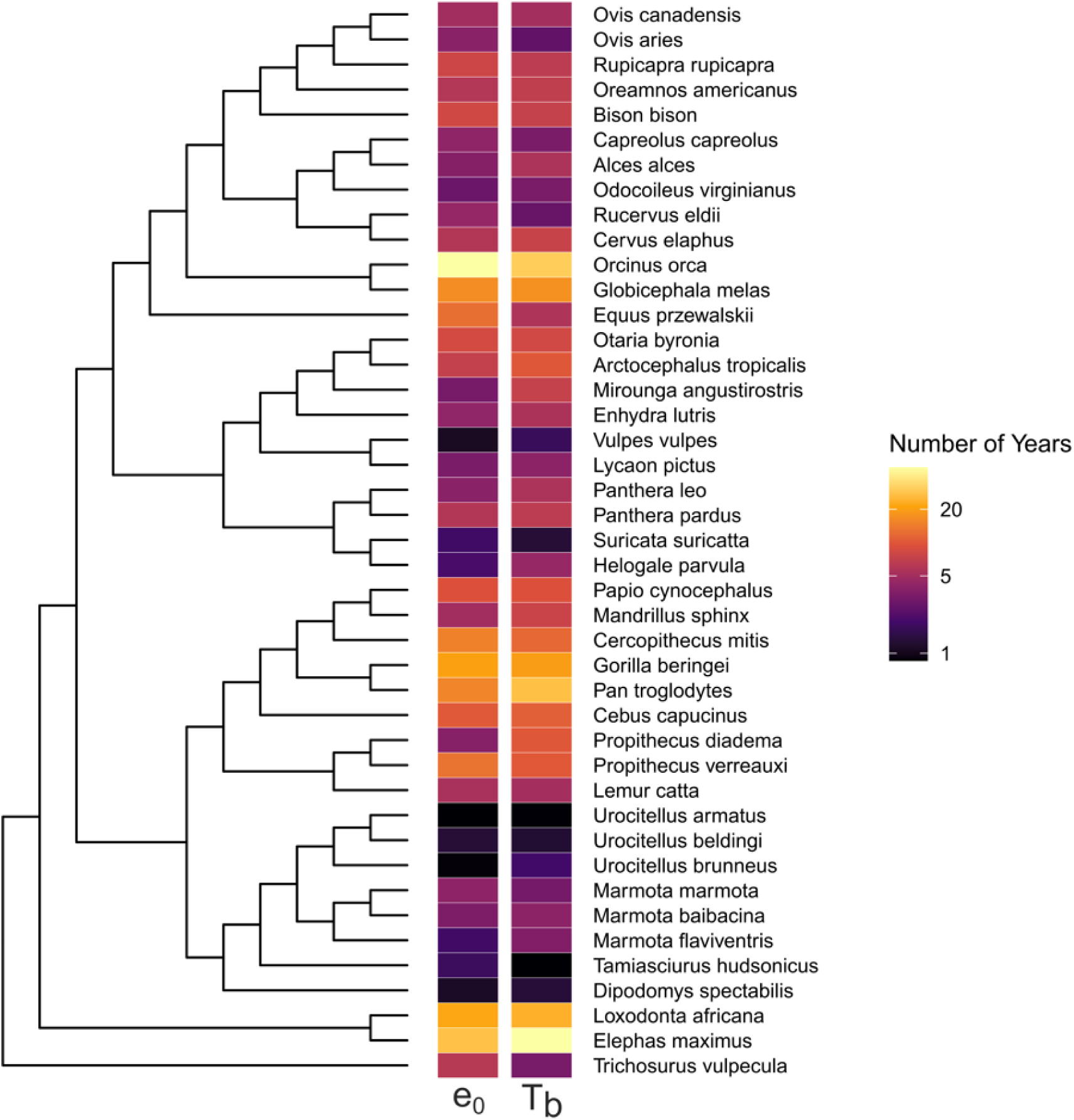
Phylogeny of all species included in the life table dataset. Each colored square represents the values of life expectancy at birth (e_0_) and generation time (T_b_) from inner to outer circles, respectively. The phylogeny was built using the “rotl” R package (Michonneau et al., 2016).

## 4) Future of malddaba

The current version of malddaba contains age-specific survival data for both sexes and age-specific reproductive data mostly on females. This content will be developed in the next coming months to further extend the range of questions that can be addressed with malddaba. Beyond the recording of our back-log catalogue on age-specific survival probabilities for many populations, one of the first logical step will be the addition of age-specific data on male reproductive success. While such data are, at least at first glance less reported in males than in females (Archer et al., 2022), there are in fact available in the literature for multiple mammalian species and multiple traits (e.g. mating success, Raveh et al., 2010; litter size, Thorley et al., 2020).

One of our major aim is also to progressively incorporate age-specific data on functional traits (e.g. morphological and physiological traits) on both males and females into malddaba. While the relevance of such traits in the context of environmental perturbations, such as climate change, is increasingly documented (Ozgul et al., 2010; Plard et al., 2015), large scale comparative analyses seeking to identify the ecological drivers of the age-specific changes in functional traits remain extremely rare. Yet, numerous empirical studies have already provided evidence that the allocation to functional traits (e.g. body mass, size of secondary sexual traits such as antler size in cervids) change over the life course and are subjected to senescence (Cambreling et al., 2023 for antler size; Nussey et al., 2011 for body mass; Cheynel et al., 2017 for immune parameters). Overall the clear influence of age-specific decline in functional traits on evolutionary dynamics (*e.g.* Bonduriansky et al., 2008 for sexual traits) combined with the increased availability of such data in the literature calls for this information to be added to malddaba.

Finally, adding age-specific data on functional traits into malddaba will offer new opportunities to investigate differences in life history strategies during the growth period across mammals. This could be done through the analyses of age-specific changes in body mass in early life. Indeed, growth trajectories are known to be extremely variable across mammals in terms of both shape and intensity (Gaillard et al., 1997), which can further impact population dynamics (Ozgul et al., 2010). In addition, this focus on early life could also take advantage of the age-specific mortality data included in malddaba. So far, the few studies performed with these data have focused on the adult or senescent stages, while comparative analyses of the age-specific changes in mortality risk in early life would be particularly interesting to better understand the usually large variation in juvenile mortality of most mammalian species (Clutton-Brock et al., 1985).

## Glossary

l_x_ series or cumulative survival: proportion of individuals born in a population that survives until age *x*.
m_x_ series or age-specific fecundity: average number of female offspring born to an individual of age *x*.
Generation time, T: representing the weighted mean age of mothers in the population.
Net reproductive rate, R0: the average number of female offspring alive at birth produced by an individual during its lifetime.
Age-specific reproductive values, V_x_: average number of remaining offspring to be produced for an individual of age *x*, standardized so *V_x_* is 1 at age 0.
Life expectancy, e_x_: average number of remaining years before death for an individual of age *x*
D_x_ series: distribution of the ages at death in a population.
S_x_ series or survival probabilities: probability for an individual alive at age *x* to survive to age *x+1*.
Longitudinal monitoring: type of monitoring in which individuals are followed through their lifetime mostly through CMR sampling.
Transversal or cross-sectional monitoring: type of monitoring in which individuals in a population are only sampled at a specific time point and not followed, which allows estimating the age distribution of alive or dead individuals for that specific date.
Population projection model (PPM): a set of demographic models that aims to project population structures from one time period to the next one, in order to predict future population changes based on current demographic transitions.

## Acknowledgments

The compilation of age-specific demographic data of vertebrates in the Laboratory of Biometry and Evolutionary Biology (LBBE) takes us back to 1985 and the research program ATIPE CNRS granted to Jean-Dominique Lebreton. We are grateful to all colleagues and students who have contributed to this data collection since then, through various research projects on comparative demography. In particular, we warmly thank Jean-Dominique Lebreton, Jean Clobert, Dominique Pontier, Jacques Trouvilliez, Dominique Allainé, Christophe Pélabon, Anne Loison, Carole Toïgo, Christophe Bonenfant, Marlène Gamelon, Vérane Berger, Morgane Tidière and Solène Cambreling. We thank Jacques Masson for creating the website of malddaba and Fabrice Hibert for designing the logo of the database. We would also like to thank all researchers, field assistants and students who participated in the monitoring and data collection efforts associated with all the mammalian populations included in malddaba.

## Author contribution

V.R., JM.G. and JF.L. collectively conceived and collected the information for this database. V.R., L.H. and B.S. built the architecture of the database. V.R., JM.G. and JF.L. wrote conjointly a first draft of this manuscript which was then commented and accepted by all authors.

## Data availability statement

All data used in this publication are available on the website of malddaba (https://malddaba.univ-lyon1.fr/).

## Supplementary information

**Appendix S1: Calculation of demographic metrics from life tables and population projection matrices**

**Appendix S2: Life table data**

