## Appendix S1 for "A MammaLian Demographic Database for comparative analyses of evolutionary biodemography: malddaba"

**Appendix S1: Calculation of demographic metrics from life tables and population projection matrices**

We will use an example using a longitudinal monitoring on wild dogs (*Lycaon pictus*) in Selous National Park (Creel and Creel 2002) to show that using either a life table, a pre-breeding census Population Projection Matrix, or a post-breeding census Population Projection Matrix allow us to compute the same demographic metrics (R code is available at <https://github.com/vronget/Malddaba/blob/main/Script_demographic_metrics.R> to compute all metrics in this appendix for this example)

**1) Life table approach**

The age-specific survival (*S_x_* series) and fecundities (*m_x_* series) are downloaded and tabulated in the life table. The *l_x_* series (cumulative survival to age *x*) is directly calculated from the *S_x_* series:

| Age | *l_x_* | *m_x_* | S_x_ |
| --- | --- | --- | --- |
| 0 | 1 | 0.000 | 0.75 |
| 1 | 0.750 | 0.000 | 0.84 |
| 2 | 0.630 | 0.070 | 0.79 |
| 3 | 0.497 | 0.210 | 0.75 |
| 4 | 0.373 | 0.440 | 0.78 |
| 5 | 0.291 | 0.840 | 0.46 |
| 6 | 0.134 | 3.000 | 0.69 |
| 7 | 0.092 | 1.610 | 0.22 |
| 8 | 0.020 | 2.500 | 1.00 |
| 9 | 0.020 | 2.000 | 1.00 |
| 10 | 0.020 | 0.000 | 0.00 |
| 11 | 0 | 0.000 | 0.00 |

From this information, one can easily calculate the net reproductive rate, *R0*, as:

$$R0=\sum_{0}^{\infty} l_{x}m_{x}$$

*R0* is equal to 1.19949 meaning that a female produces on average about 1.20 daughters throughout its lifetime.

We can calculate the cohort generation time (*T_c_*) as:

$$Tc=\frac{\sum_{0}^{\infty} {xl}_{x}m_{x}}{R0}$$

*T_c_* is equal to 5.424, meaning that the mean age of reproduction by a cohort is about 5 years and 5 months.

The growth rate *r* can be calculated by numerically solving the Euler-Lotka equation (see R code for the numerical solving).

$$\sum_{0}^{\infty} {e^{-rx}l}_{x}m_{x}=1$$

We estimated here *r* to be 0.0338 and the natural rate of increase (*λ*) can be calculated as $e^{r}=1.034$

We can also calculate the generation time after discounting the influence of population growth, which corresponds to the *T_b_* metric proposed by Leslie (1966) as:

$$Tb=\frac{\sum_{0}^{\infty} {xe^{-rx}l}_{x}m_{x}}{\sum_{0}^{\infty} {e^{-rx}l}_{x}m_{x}}$$

*T_b_* is equal to 5.343, meaning that the weighted mean age of mothers right before giving birth is about 5 years and 4 months. In a stationary population with a constant population size over time (i.e*. r* = 0), *T_b_* and *T_c_* are identical.

**1) Matrix population projection matrix approach**

With the pre-breeding census formulation, the recruitment of the population is measured when offspring are 1 year of age. The age-specific values thus correspond to the number of female offspring produced by a mother at a given age (*m_x_*) times the offspring survival between birth and 1 year of age (first row of the matrix) while the survival transition from one age to the next one are displayed in the diagonal of the matrix.

*Pre-breeding census Population Projection Matrix*

|  | *1yr* | *2yr* | *3yr* | *4yr* | *5yr* | *6yr* | *7yr* | *8yr* | *9yr* | *10yr* |
| --- | --- | --- | --- | --- | --- | --- | --- | --- | --- | --- |
| *1yr* | 0 | 0.0525 | 0.1575 | 0.33 | 0.63 | 2.25 | 1.2075 | 1.875 | 1.5 | 0 |
| *2yr* | 0.84 | 0 | 0 | 0 | 0 | 0 | 0 | 0 | 0 | 0 |
| *3yr* | 0 | 0.79 | 0 | 0 | 0 | 0 | 0 | 0 | 0 | 0 |
| *4yr* | 0 | 0 | 0.75 | 0 | 0 | 0 | 0 | 0 | 0 | 0 |
| *5yr* | 0 | 0 | 0 | 0.78 | 0 | 0 | 0 | 0 | 0 | 0 |
| *6yr* | 0 | 0 | 0 | 0 | 0.46 | 0 | 0 | 0 | 0 | 0 |
| *7yr* | 0 | 0 | 0 | 0 | 0 | 0.69 | 0 | 0 | 0 | 0 |
| *8yr* | 0 | 0 | 0 | 0 | 0 | 0 | 0.22 | 0 | 0 | 0 |
| *9yr* | 0 | 0 | 0 | 0 | 0 | 0 | 0 | 1.00 | 0 | 0 |
| *10yr* | 0 | 0 | 0 | 0 | 0 | 0 | 0 | 0 | 1.00 | 0 |

The natural rate of increase (*λ*) corresponding to the largest eigenvalue of this matrix is 1.034364 and *r* can be calculated as log(*λ*) We get a value of 0.0338 for *r*, the same as for the life table approach.

The generation time can be calculated as the inverse of the overall elasticity to recruitment (Bienvenu and Legendre 2015). The elasticity matrix E is calculated using the following formula:

$$E_{ij}=\frac{d\lambda}{da_{ij}}\times\frac{a_{ij}}{\lambda}$$

with *λ* the natural rate of increase and *a_ij_* the transition rate from age i to j.

*Matrix of elasticities*

|  | *1yr* | *2yr* | *3yr* | *4yr* | *5yr* | *6yr* | *7yr* | *8yr* | *9yr* | *10yr* |
| --- | --- | --- | --- | --- | --- | --- | --- | --- | --- | --- |
| *1yr* | 0 | 0.0077 | 0.0177 | 0.0269 | 0.0387 | 0.0614 | 0.0220 | 0.0073 | 0.0056 | 0 |
| *2yr* | 0.1871 | 0 | 0 | 0 | 0 | 0 | 0 | 0 | 0 | 0 |
| *3yr* | 0 | 0.1794 | 0 | 0 | 0 | 0 | 0 | 0 | 0 | 0 |
| *4yr* | 0 | 0 | 0.1618 | 0 | 0 | 0 | 0 | 0 | 0 | 0 |
| *5yr* | 0 | 0 | 0 | 0.1349 | 0 | 0 | 0 | 0 | 0 | 0 |
| *6yr* | 0 | 0 | 0 | 0 | 0.0963 | 0 | 0 | 0 | 0 | 0 |
| *7yr* | 0 | 0 | 0 | 0 | 0 | 0.0349 | 0 | 0 | 0 | 0 |
| *8yr* | 0 | 0 | 0 | 0 | 0 | 0 | 0.0129 | 0 | 0 | 0 |
| *9yr* | 0 | 0 | 0 | 0 | 0 | 0 | 0 | 0.0056 | 0 | 0 |
| *10yr* | 0 | 0 | 0 | 0 | 0 | 0 | 0 | 0 | 0 | 0 |

The overall elasticity to recruitment, which corresponds to the inverse of the sum of the elements of the first row of the elasticity matrix, equals 5.343, meaning that the weighted mean age of females at parturition (i.e. the generation time, *T_b_*) is about 5.34 years. Note that this value perfectly matches the estimation of *T_b_* based on the life table approach.

*Post-breeding census Population Projection Matrix*

|  | *0yr* | *1yr* | *2yr* | *3yr* | *4yr* | *5yr* | *6yr* | *7yr* | *8yr* | *9yr* | *10yr* |
| --- | --- | --- | --- | --- | --- | --- | --- | --- | --- | --- | --- |
| *0yr* | 0 | 0.0588 | 0.1659 | 0.33 | 0.6552 | 1.38 | 1.1109 | 0.55 | 2 | 0 | 0 |
| *1yr* | 0.75 | 0 | 0 | 0 | 0 | 0 | 0 | 0 | 0 | 0 | 0 |
| *2yr* | 0 | 0.84 | 0 | 0 | 0 | 0 | 0 | 0 | 0 | 0 | 0 |
| *3yr* | 0 | 0 | 0.84 | 0 | 0 | 0 | 0 | 0 | 0 | 0 | 0 |
| *4yr* | 0 | 0 | 0 | 0.75 | 0 | 0 | 0 | 0 | 0 | 0 | 0 |
| *5yr* | 0 | 0 | 0 | 0 | 0.78 | 0 | 0 | 0 | 0 | 0 | 0 |
| *6yr* | 0 | 0 | 0 | 0 | 0 | 0.46 | 0 | 0 | 0 | 0 | 0 |
| *7yr* | 0 | 0 | 0 | 0 | 0 | 0 | 0.69 | 0 | 0 | 0 | 0 |
| *8yr* | 0 | 0 | 0 | 0 | 0 | 0 | 0 | 0.22 | 0 | 0 | 0 |
| *9yr* | 0 | 0 | 0 | 0 | 0 | 0 | 0 | 0 | 1.00 | 0 | 0 |
| *10yr* | 0 | 0 | 0 | 0 | 0 | 0 | 0 | 0 | 0 | 1.00 | 0 |

With the post-breeding census formulation, the recruitment of the population is measured when offspring are just born. The reproduction values thus correspond to the number of female offspring produced by a mother at a given age (*m_x_*) times the mother survival between the previous census and the current one.

The natural rate of increase (*λ*) corresponding to the largest eigenvalue of this matrix is 1.034. Note that this value is exactly the same as that estimated from the pre-breeding census model. Any difference between pre- and post-breeding census models come from errors when identifying the recruitment entries of the matrix (Kendall et al. 2019). Martin et al. (2025) recently provided a new R-package that allows building rightly any pre- or post-breeding census matrix and thereby avoiding any error in population growth estimate.
