## Appendix S2 for "A MammaLian Demographic Database for comparative analyses of evolutionary biodemography: malddaba"

**Appendix S2: Life table data**

**1) Protocol of Life Table Construction**

We compiled malddaba records to build female life tables for different populations. For a population to be eligible for life table construction, both the *S_x_* series (survival transitions from birth to the last age) and the *m_x_* series (reproduction transitions from the age at first reproduction to the last age) for female specifically and for the same population were required. We began each time by first building the complete age-dependent *S_x_* series, followed by the associated age-dependent *m_x_* series (all information about the construction of the specific series is reported in the malddaba life table dataset in Rdata format available at <https://malddaba.univ-lyon1.fr/pages/life_table_data.php>, specifically in the data_used table of the Rdata; see table S1 for an explanation of each column associated with the life table data file).

*S_x_ series construction*

*S_x_* series were constructed differently depending on the type of survival data available:

- For longitudinal survival data *S_x_* was directly reported from the *S_x_* series in the malddaba record.

- For transversal data associated with the *d_x_* series (when only age at death is reported), we considered that the distribution of ages at death matched the *d_x_* series of the population and calculated *S_x_* using the following formula $S_{x}=\frac{d_{x+1}}{d_{x}}$. When the original publication reported an age-dependent model to smooth the distribution of ages at death, we preferred to compute the *S_x_* based on this smoothed distribution.

- For transversal data associated with *l_x_* series (when the age of alive sampled individual is reported), we considered that the distribution of ages at death matched the *l_x_* series of the population and calculated the *S_x_* series based on the following formula $S_{x}=\frac{l_{x+1}}{l_{x}}$. When the original publication reported an age model to smooth the distribution of alive individuals, we chose to compute the *S_x_* based on this smoothed distribution.

We verified that the *S_x_* series started from birth (age 0). Sometimes, juvenile and adult survival rates were reported in different malddaba records because they originated from different publications. In that case, we maintained our inclusion criterion, which required that both juvenile and adult survival data come from the same population.

For a large proportion of the *S_x_* series, the survival probability at the last age was not equal to 0, meaning that the *S_x_* series was right-censored due to the deaths of the few oldest individuals not being reported. In that case, we checked the last *l_x_* at the censored age; if the value was lower than 5%, we chose to artificially end the *S_x_* series by adding a survival probability of 0 at the next age. If the last *l_x_* value was higher than 5%, we either did not consider that population if too many individuals were censored, or if the series was higher but close to the 5% threshold, we extended the *S_x_* series by adding a survival probability equal to that of the last age observed until the *l_x_* was lower than 5%. We then added a survival probability of zero to finish the *S_x_* series. All decisions made to construct the *l_x_* series are either reported as categories or noted in the comment column if specific transformations were necessary.

*m_x_ series construction*

The construction of the *m_x_* series also depended on the type of reproduction data available in malddaba:

- Sometimes *m_x_* was directly calculated in the original record.

- When the number of offspring produced per age was available, we simply divided it by 2 to obtain the average number of daughters, assuming that the sex ratio was balanced between male and female offspring produced.

- For monotocous species (*i.e.* litter size of 1) when the probability to reproduce was available, we simply divided this probability by 2 considering again a balanced sex ratio to get the *m_x_* series.

-for polytocous species (*i.e.* litter size > 1), both the probability to reproduce and the litter size per age were necessary if the *m_x_* or the number of offspring per age was not available. In that case, we calculated the *m_x_* series by multiplying the probability of reproducing and the average litter size per age divided by 2.

For some reproduction data, *m_x_* values were missing for very old ages because less individuals were sampled for the reproduction analysis compared to the survival analyses in this population. In that case we extended the *m_x_* series using the *m_x_* value at the last age reported in the data. As for survival, all decisions used for the construction of the *m_x_* series are reported in the dataset.

**2) Demographic outputs of life tables**

Based on the full life table (*m_x_* and *S_x_* series) built, we were able to compute several demographic outputs using the following formulas:

- *l_x_* series (proportion of individuals still alive at age *x*)

$$l_{x+1}=l_{x}S_{x} with l_{0}=1$$

- *d_x_* series (proportion of individuals dying at age *x*)

$$d_{x}=l_{x+1}-l_{x}$$

- *e_x_* series, life expectancy at age x (average number of remaining years of life at age *x*)

$$e_{x}=\frac{1}{l_{x}}\sum_{x}^{\infty} l_{i}-0.5$$

- *r*, the asymptotic growth rate of the population, computing by solving numerically the following Euler-Lotka equation

$$\sum_{0}^{\infty} {e^{-rx}l}_{x}m_{x}=1$$

- *R0*, the net reproductive rate of the population

$$R0=\sum_{0}^{\infty} l_{x}m_{x}$$

- *Vx* series, the reproductive values for each age (Fisher, 1930) (average number of individuals that will be produced by a female of age *x* for her remaining lifetime, standardized so *V_0_*=1)

$$V_{x}=\frac{e^{-rx}}{l_{x}}\sum_{x+1}^{\infty} e^{-ri}l_{i}m_{i}$$

*T_b_*, generation time of the population (Gaillard et al., 2005) (mean age at reproduction in years in the population accounting for the stable age distribution)

$$Tb=\frac{\sum_{0}^{\infty} {xe^{-rx}l}_{x}m_{x}}{\sum_{0}^{\infty} {e^{-rx}l}_{x}m_{x}}$$

*T_c_*, cohort generation time (Steiner et al., 2014) (mean age at reproduction in years for a specific cohort)

$$Tc=\frac{\sum_{0}^{\infty} {xl}_{x}m_{x}}{R0}$$

Table S1: Information reported in the malddaba life table dataset (See text for more details about the life table construction procedure and the formula used to calculate the demographic outputs)

| Table | Column | Description |
| --- | --- | --- |
| species | species | Species latin names (genus then species) |
| data_used |  | Information on the malddaba record used and the procedure used to compute the female life table |
|  | Immature_survival_life_table Id | Identifier of the malddaba record used to compute immature survival probabilities |
|  | Adult_survival_life_table_id | Identifier of the malddaba record used to compute adult survival probabilities |
|  | Censored_survival | Whether survival series was artificially ended by censoring with a null survival at last age |
|  | Extended_survival | Whether a procedure to extend survival probability at old ages was applied |
|  | Survival_modeled | Whether survival from reported age-dependent model was used instead of the raw full age estimates |
|  | Survival_comments | Any comments associated with a specific procedure used to compute *S_x_* series for this population |
|  | mx_id | Identifier of the malddaba record used to compute *m_x_* series if *m_x_* was directly available |
|  | N_offspring | Identifier of the malddaba record used to compute *m_x_* series using the number of offspring produced per age |
|  | P_reproduce_id | Identifier of the malddaba record used to compute *m_x_* series using the probability to reproduce per age |
|  | Litter_size_id | Identifier of the malddaba record used to compute *m_x_* series using the average litter size per age |
|  | Censored_reproduction | Whether *m_x_* series was artificially censored at last age based on the *S_x_* series |
|  | Extended_reproduction | Whether a procedure to extend *m_x_* series at old ages was applied |
|  | Reproduction_modeled | Whether reproduction from reported age-dependent model was used instead of the raw full age estimates |
|  | Reproduction_comments | Any comments associated with a specific procedure used to compute reproduction for this population |
|  | Age_interval | Age interval in years used for the building of the life table |
| Life_table |  | Raw life table computed |
|  | Age | Age in years |
|  | Sx | Age-specific *S_x_* values (survival probabilities) |
|  | lx | Age-specific *l_x_* values (cumulative survival) |
|  | mx | Age-specific *m_x_* values (average number of daughters produced per females) |
| Life_table_extended |  | Life table with additional columns computed based on demographic outputs |
|  | Age | Age in years |
|  | Sx | Age-specific *S_x_* values (survival probabilities) |
|  | lx | Age-specific *l_x_* values (cumulative survival) |
|  | mx | Age-specific *m_x_* values (average number of daughters produced per females) |
|  | dx | Age-specific *d_x_* values |
|  | ex | Age-specific life expectancies |
|  | Vx | Age-specific reproductive values |
| Demographic_metrics |  | Demographic metrics computed to describe the dynamic of the population |
|  | AFR | Age at first reproduction in years |
|  | R0 | Net reproductive rate |
|  | r | Asymptotic growth rate |
|  | Tb | Generation time corrected by the age structure of the population in years |
|  | Tc | Cohort generation time in years |
